# Near-future warming amplifies natural heatwave impacts and reorganizes freshwater communities

**DOI:** 10.64898/2026.07.01.735828

**Authors:** Sara Nouere, Martin Schäfer, Guangqiong Li, Martin Lohr, Dieter Ebert, Shuqing Xu

## Abstract

Future climate change may reshape ecological communities not only by increasing mean temperature, but also by altering the consequences of increasingly frequent heatwaves. Predicting these effects requires understanding how background warming interacts with short heatwaves in natural communities, where responses can arise through direct thermal stress and species interactions. We tested this using 32 outdoor freshwater mesocosms exposed to sustained near-future warming while capturing a documented natural heatwave. Warming raised temperature maxima that exceeded the thermal threshold of the pond snail, a main grazer in the community. Warmed communities showed lower grazer abundance, increased macrophyte and insect herbivore abundance, reduced phytoplankton biomass, and lower zooplankton density. Complementary assays showed that heatwave-level temperatures promoted macrophyte growth and reduced grazer survival, whereas reduced zooplankton performance mainly reflected indirect warming effects via food-web cascades. Thus, near-future warming can amplify natural heatwave impacts by exceeding consumer thermal thresholds and propagating through species interactions.

## Introduction

Climate warming is reshaping ecological communities by altering both the performance of individual species and the interactions that regulate population dynamics, coexistence, and ecosystem functioning (Parmesan & Yohe 2003; Kearney *et al*. 2009; Li *et al*. 2011; Paull & Johnson 2014). Because temperature influences metabolism, feeding, growth, and thus demography, its effects can propagate through food webs and generate community-level outcomes that cannot be predicted from single-species responses alone (Carpenter *et al*. 1985; Tylianakis *et al*. 2008; Kearney *et al*. 2009). Freshwater ecosystems are particularly sensitive to climatic change because they respond rapidly to physical forcing and often show strong trophic coupling between producers and consumers (Adrian *et al*. 2009).

Extreme heat events are increasing in frequency and intensity and can drive abrupt ecological change when organisms are exposed to temperatures near or beyond their physiological limits (Frölicher & Laufkötter 2018; Stillman 2019; Perkins-Kirkpatrick & Lewis 2020). In freshwater systems, recent work has highlighted the importance of subsurface heat extremes and the loss of vertical thermal refuges, which can strongly shape exposure risk across communities (Woolway *et al*. 2021). Yet experimental tests at the community level remain limited. Heatwaves are difficult to predict and capture, and many warming experiments still focus primarily on changes in mean temperature, thereby underrepresenting the temporal variability and short-lived extremes that may be most consequential when thermal thresholds are crossed (Baum *et al*. 2026).

A major reason for this gap is methodological. Outdoor warming experiments at ecologically relevant scales are often technically demanding, energy-intensive, and difficult to replicate, which limits both accessibility and statistical power (Yvon-Durocher *et al*. 2010; Stewart *et al*. 2013). At the same time, theoretical and empirical work increasingly emphasizes that climate-change experiments should incorporate not only changes in mean temperature, but also changes in variability and extremes, because biological responses to temperature are nonlinear (Thompson *et al*. 2013; Vasseur *et al*. 2014; Sinclair *et al*. 2016). Cost-effective warming systems that operate at scale while preserving natural temporal variation are therefore essential for testing how climate change restructures ecological communities under realistic conditions.

Here, we developed a scalable, solar-powered warming platform for 32 outdoor freshwater mesocosms and used it to study a multitrophic community consisting of the floating duckweed *Spirodela polyrhiza*, the grazing snail *Lymnaea stagnalis*, the waterlily aphid *Rhopalosiphum nymphaeae*, phytoplankton, and the zooplankton grazer *Daphnia magna*. The system generated a near-future warming treatment while preserving natural weather-driven temperature dynamics across the growing season. Using this platform, we tested whether near-future warming amplifies the community impacts of a natural heat event by eroding thermal refuges for consumers and altering trophic interactions.

## Results

### A scalable, low-cost warming system tracked natural thermal dynamics

We established 32 outdoor mesocosms (∼ 5,400 L; 16 warmed and 16 ambient temperature controls) containing a multitrophic freshwater community and implemented a solar-powered system to elevate water temperature in the warmed treatment relative to controls (Fig. 1a,b). One-time infrastructure costs were approximately €1,850 per pond, and most components were reusable (Table S1). Operational electricity demand was driven mainly by the circulation pumps (∼20 W per pond), resulting in low annual energy costs of approximately €45 per pond across the 32-pond platform (Table S2).

**Figure 1.**
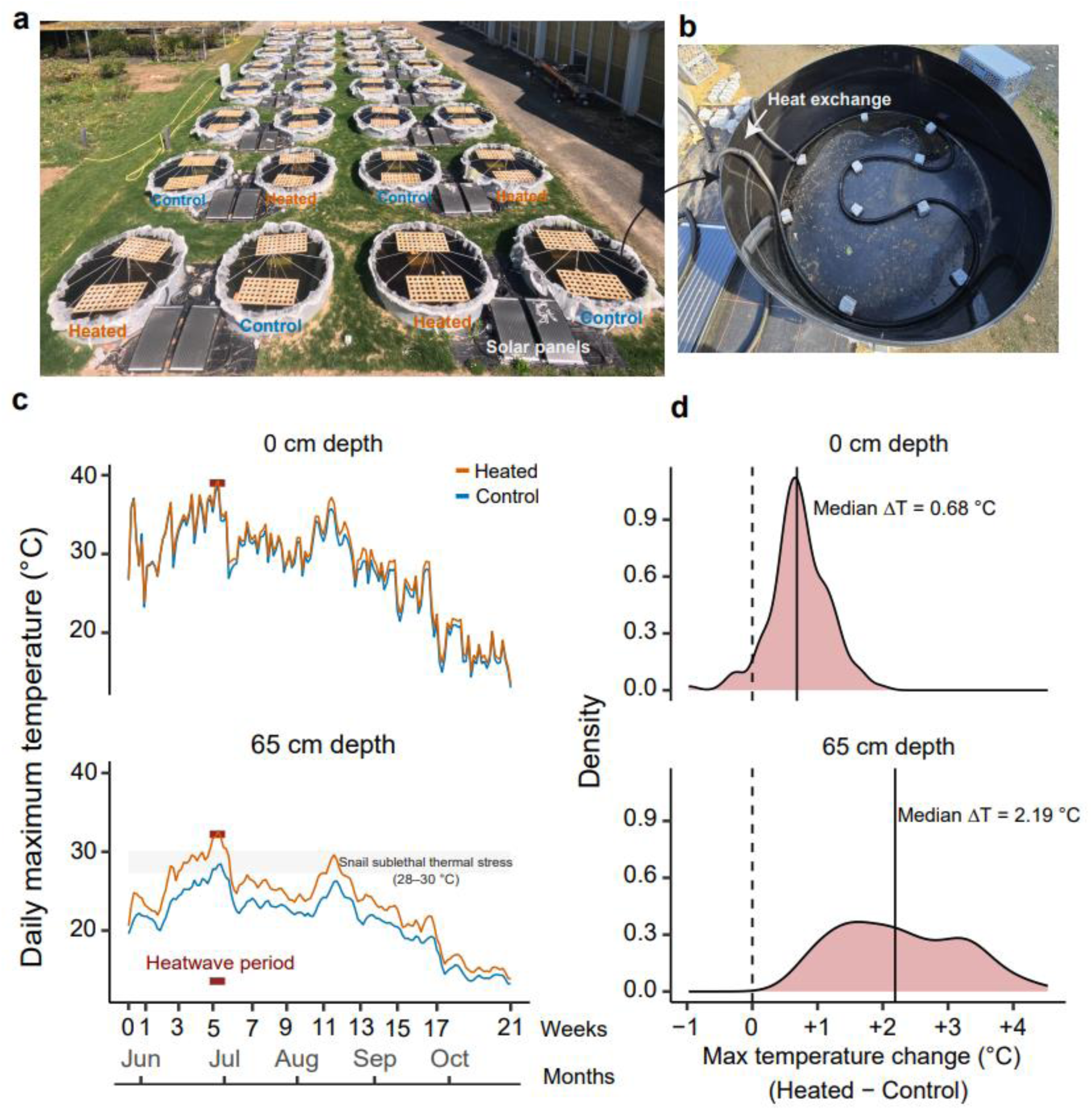
Warming amplified a regional heatwave beyond the sublethal thermal stress threshold of snails *Lymnaea stagnalis*, eliminating deep-water refuges. **a,** Overview of the outdoor mesocosm experiment showing heated (n = 16; orange) and control (n = 16; blue) ponds and solar-powered warming infrastructure. **b,** Example of the internal heat-exchange system used to elevate water temperature in heated ponds. **c,** Daily maximum temperatures in heated (orange) and control (blue) ponds at 0 and 65 cm depth across the experimental period. The red band indicates the documented regional heatwave (Mainz, Western Germany, 28 June–02 July 2025). The grey shaded area refers to the sublethal thermal range of *L. stagnalis* (28–30 °C). During the heatwave, elevated temperature drove deep-water temperatures in heated ponds beyond this range, whereas control ponds retained sublethal conditions at depth. **d,** Distribution of daily treatment differences in maximum temperatures (ΔT = heated – control) for each depth. Vertical solid lines indicate median ΔT and dashed lines indicate zero difference. Positive values indicate days when heated ponds were warmer than controls.

Continuous temperature logging at the surface and at 65 cm depth showed that warmed ponds maintained consistently higher mean temperatures than controls (median ΔT: 1.24 °C at the surface, 1.91 °C at depth; Fig. S1a,b) while preserving natural temporal variation. During the regional heatwave in early July 2025, maximum temperatures increased in both treatments, but warmed ponds reached higher maxima than controls (Fig. 1c,d). At 65 cm depth, the median warmed–control difference in daily maximum temperature was 2.19 °C (Fig. 1d), consistent with a modest near-future warming scenario (IPCC 2018). During this period, temperatures at 65 cm depth in warmed ponds exceeded the reported sublethal thermal stress threshold of *Lymnaea stagnalis* (30 °C; Fig. 1c) (Hoefnagel & Verberk 2017), whereas control ponds remained below this threshold.

### Warming altered community dynamics across trophic levels

To test whether warming changed community dynamics under field conditions, we quantified the seasonal trajectories of the major trophic groups in the experimental ponds. Across the growing season, warmed ponds diverged from controls in multiple components of the community (Fig. 2; Table S3). Duckweed population size increased more strongly in warmed ponds than in controls (*F_1,15_* = 14.38, *P* = 0.001; Fig. 2a), which appeared shortly after the heatwave. In addition, fronds collected from heated ponds were both heavier (*P* = 0.05) and larger in area (*P* = 0.001; Fig. S3), indicating enhanced plant performance under warming. In contrast, the grazing snail *L. stagnalis* declined strongly in heated ponds (*F*_1,15_ = 32.47, *P* < 0.001; Fig. 2b). Across ponds, duckweed cover was negatively correlated with snail abundance (β = −0.10 ± 0.046, *F_1, 219_* = 4.70, *P* = 0.03), consistent with reduced grazing pressure contributing to increased plant dominance.

**Figure 2.**
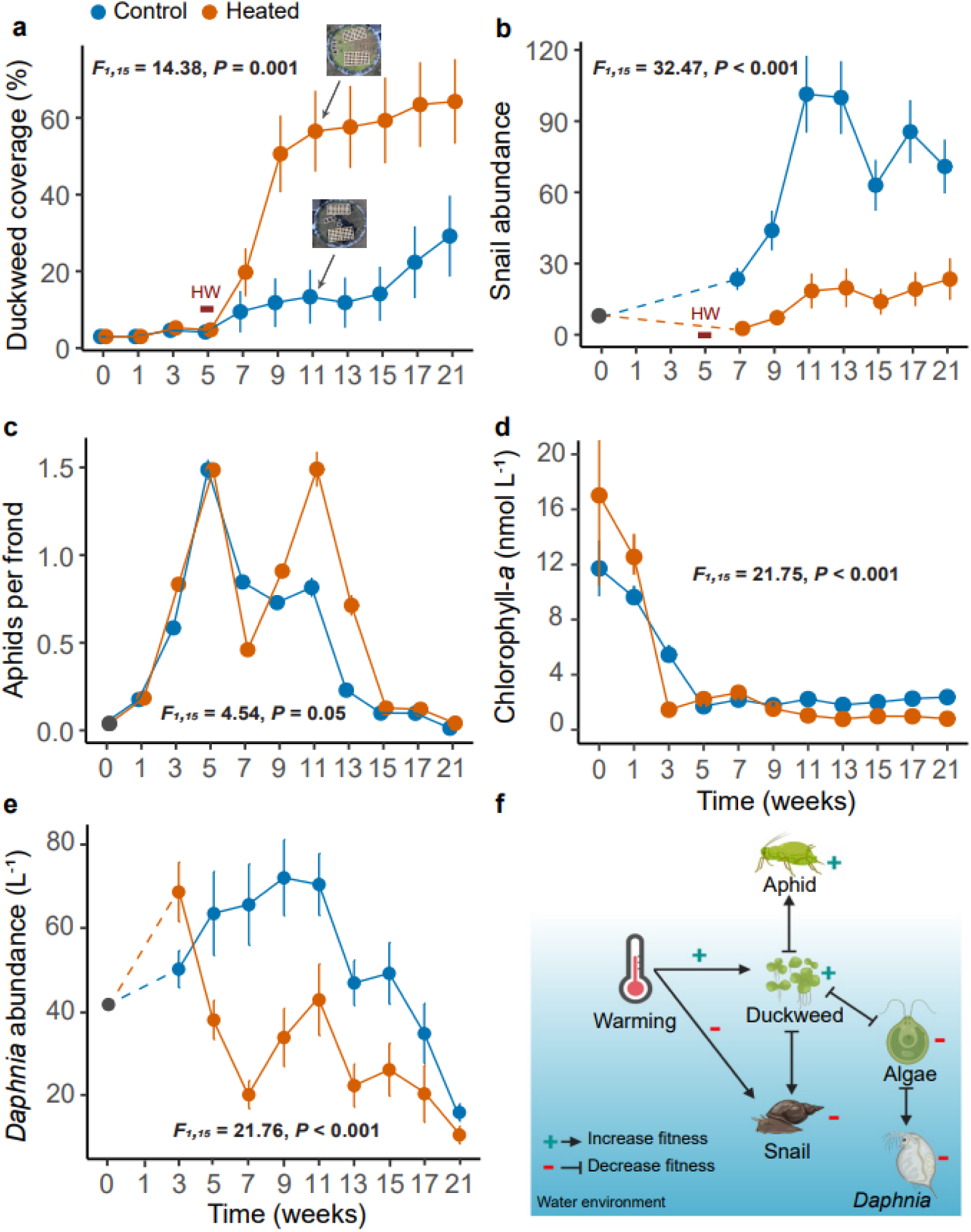
Warming altered population dynamics in experimental ponds during the 2025 growing season. **a,** Duckweed surface coverage (%) in heated (orange) and control (blue) ponds. Inset pictures show representative ponds at week 11 (top: heated; bottom: control). **b,** Snail abundance. Week 0 represents the initial total count conducted at the start of the experiment. Weeks 7–21 show snail density from a standardised vertical section of pond wall (representing 1/12 of the total pond wall). **c,** Aphid density (aphids per frond). **d,** Chlorophyll-*a* concentrations. **e,** *Daphnia* abundance. **f,** Simplified species interaction network of the experimental community under warming. Species icons in panel f were created with BioRender.com. Red horizontal bar indicates the heatwave (HW) period (28 June–02 July 2025). Points represent means (n = 16 per treatment), and error bars indicate ± SE. Population dynamics were analysed using linear mixed-effects models including treatment and time point as fixed effects, and pond and block as random effects.

Changes under warming extended to other trophic levels. Aphid density per frond was higher in warmed ponds (*F*_1,15_ = 4.54, *P* = 0.05; Fig. 2c), and at the community level, total aphid abundance (density scaled by duckweed coverage) was significantly elevated under warming (β = 0.20 ± 0.12, *F*_1,15_ = 11.33, *P* = 0.004). At the same time, phytoplankton levels based on chlorophyll-*a* concentrations were lower under warming (*F*_1,15_ = 21.75, *P* < 0.001; Fig. 2d), and *Daphnia* abundance also declined (*F*_1,15_ = 21.76, *P* < 0.001; Fig. 2e). Overall, warming was associated with a shift toward greater duckweed dominance, reduced snail abundance, increased aphid populations, lower phytoplankton biomass, and reduced *Daphnia* abundance.

### Heatwave temperature increased duckweed growth but reduced herbivory

As the interactions between the duckweed and snail appeared central to the community, we next tested how temperatures spanning the range observed in the ponds affected duckweed growth and snail herbivory in controlled climate-chamber assays (20, 24, 28, and 32 °C; Fig. 3). Duckweed growth increased strongly across the thermal gradient (*F*_3,30_ = 39.72, *P* < 0.001; Fig. 3a), reaching its highest value at 32 °C. In contrast, snail consumption showed a nonlinear response to temperature (Fig. 3a). Consumption was highest at 28 °C and then declined sharply at 32 °C. Snail mortality also increased at the highest temperature, with no mortality at 20 or 24 °C and pronounced mortality at 32 °C (Fig. 3b). These results show that temperatures reached during the heatwave can simultaneously promote duckweed growth while reducing snail feeding and survival.

**Figure 3.**
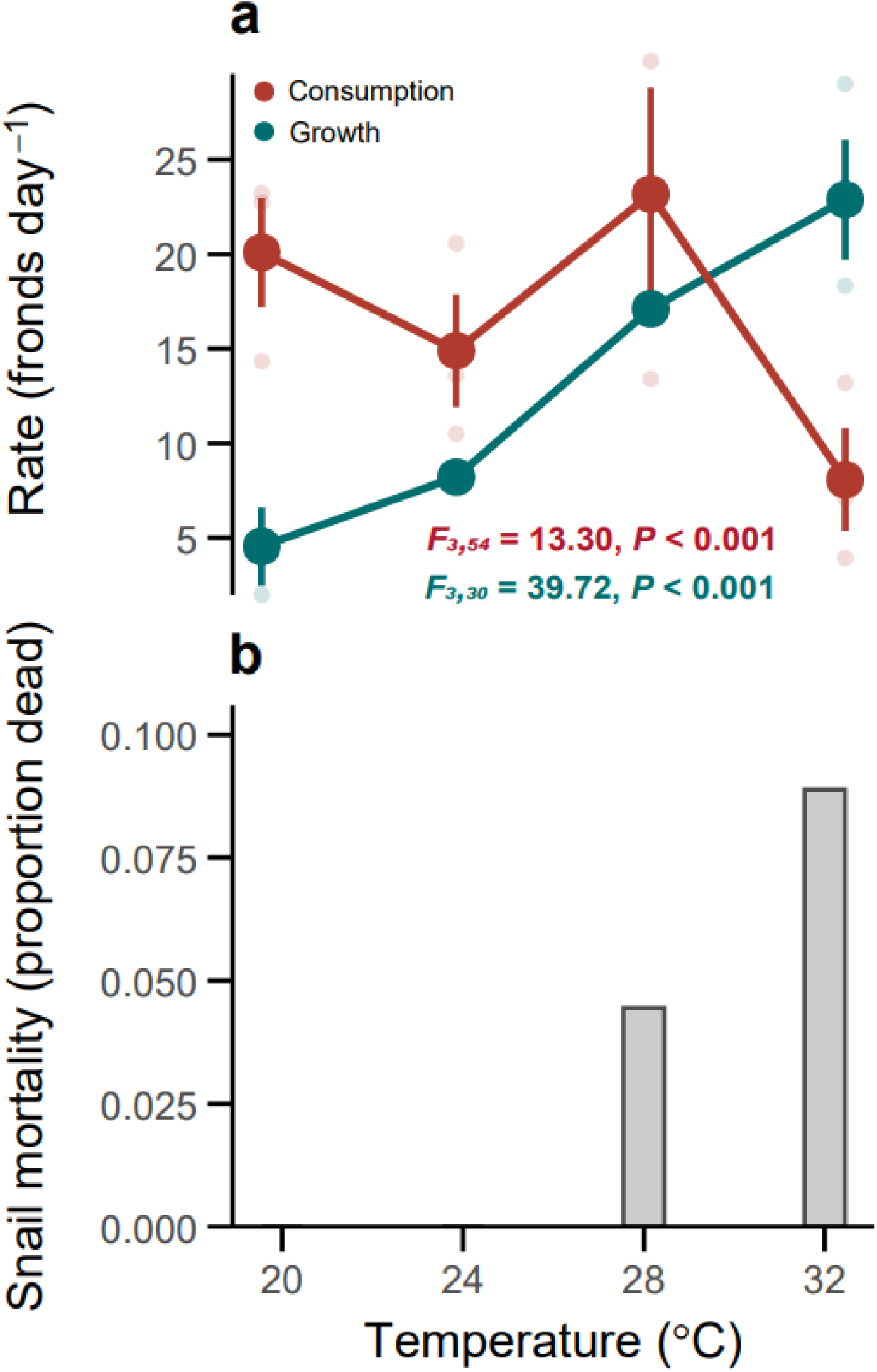
Elevated temperature directly alters plant growth, snail consumption, and snail mortality. **a,** Plant growth (green) and snail consumption (red) across four experimental temperature treatments (20, 24, 28, and 32 °C). Growth rates were measured in control microcosms without snails, whereas consumption was quantified in herbivory treatments containing three *L. stagnalis* individuals. Points represent means from three independent experimental rounds, and error bars indicate standard error (SE) across rounds. Faint points refer to round-level means. Panel **a** reports results from linear mixed-effects models with temperature as a fixed effect, and round as a random effect. **b,** Temperature-dependent snail mortality during the 22-hour feeding period. Bars show the mean proportion of snails that died, averaged across experimental rounds.

### Warming affected duckweed performance in the community

To test how outdoor warming influenced duckweed growth, we performed reciprocal transplant experiments using populations collected at week nine (4 weeks after the heatwave) (Fig. 4a). Populations originating from warmed ponds showed higher growth rates than populations from control ponds across both environments (*F_1,31_* = 10.42, *P* = 0.003; Fig. 4a), whereas host pond environment itself did not significantly affect growth (*F_1,15_* = 2.62, *P* = 0.13). Thus, differences among populations were stronger than the immediate effect of host environment in this assay.

**Figure 4.**
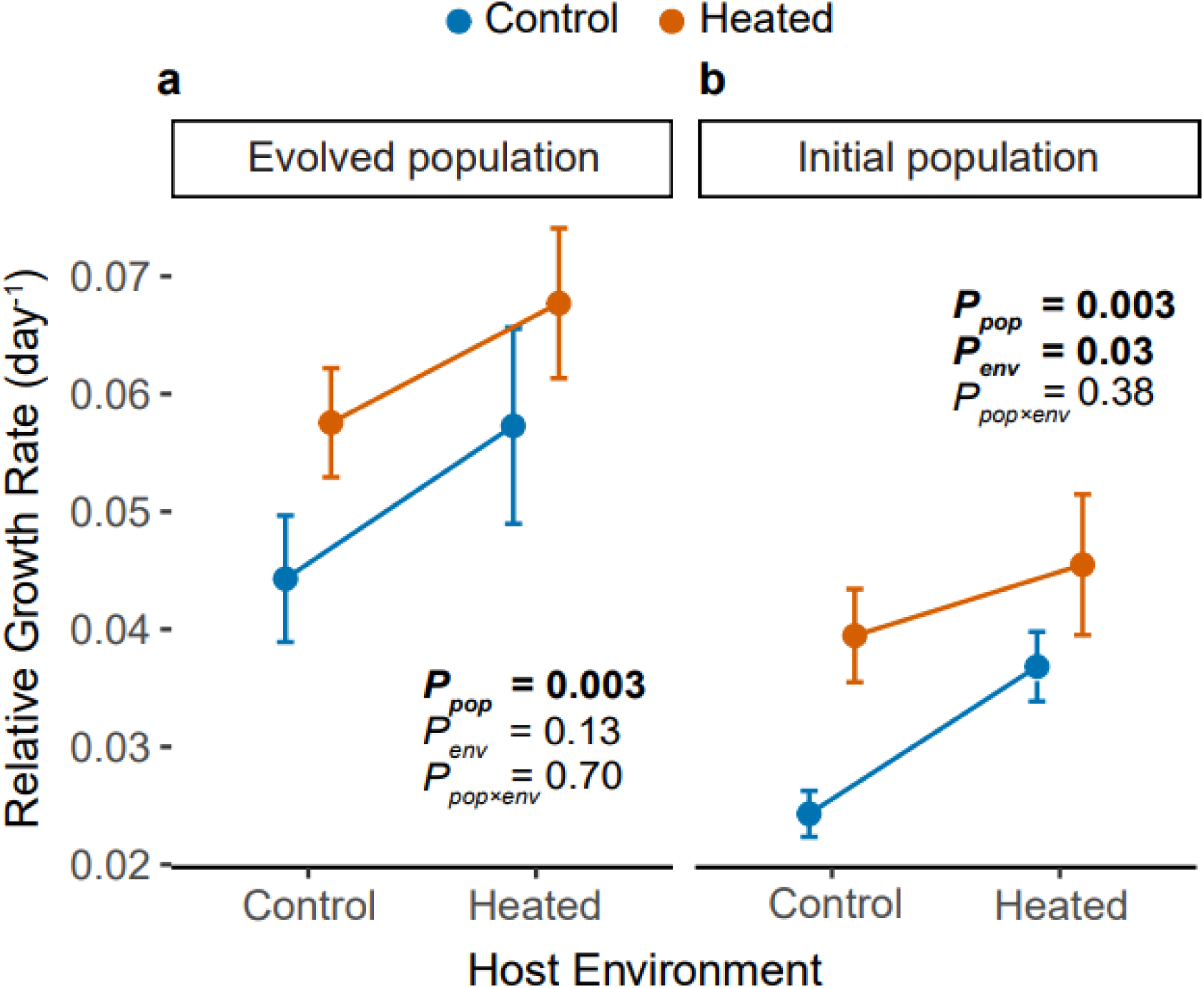
Evolutionary and plastic contributions to duckweed growth under warming. Relative growth rate (RGR) of duckweed populations in reciprocal transplant assays. **a,** RGR (day^-1^) of evolved populations (mixed genotypes after nine weeks of selection). **b,** RGR (day^-1^) of initial populations (assembled from individually maintained genotypes at equal frequencies). Populations originating from control (blue) and heated (orange) treatments were grown in control or heated host environments. Points represent mean values and error bars indicate ± SE. P-values were obtained from linear mixed-effects models including population origin and host environment as fixed effects, and pond and block as random effects.

To distinguish changes associated with changes in genotype frequencies from environmentally induced plasticity, we reconstructed the populations with the initial genotype frequency using individually maintained duckweed genotypes in both control and warmed ponds at equal frequencies and repeated the reciprocal transplant (Fig. 4b). In this assay, both host environment (*F_1,15_* = 5.64, *P* = 0.03) and genotype origin (*F_1,31_* = 10.64, *P* = 0.003) significantly affected duckweed growth. Overall, evolved populations showed significantly higher growth rates than initial populations (*F*_1,124_ = 29.92, *P* < 0.001). These data suggest that while warming itself can increase duckweed growth, the effects were likely reduced by evolutionary changes in genotype frequencies.

### Water-mediated conditions constrain *Daphnia* growth under warming

To test whether *Daphnia* performance under warming was driven by direct temperature effects and/or by warming-induced changes in pond conditions, we carried out a reciprocal transplant assay after 14 weeks of experimental warming (Fig. 5a). Individuals from warmed and control ponds were transplanted into both thermal environments using either local water or water from the alternative treatment, allowing us to separate direct thermal effects from water-mediated environmental effects.

**Figure 5.**
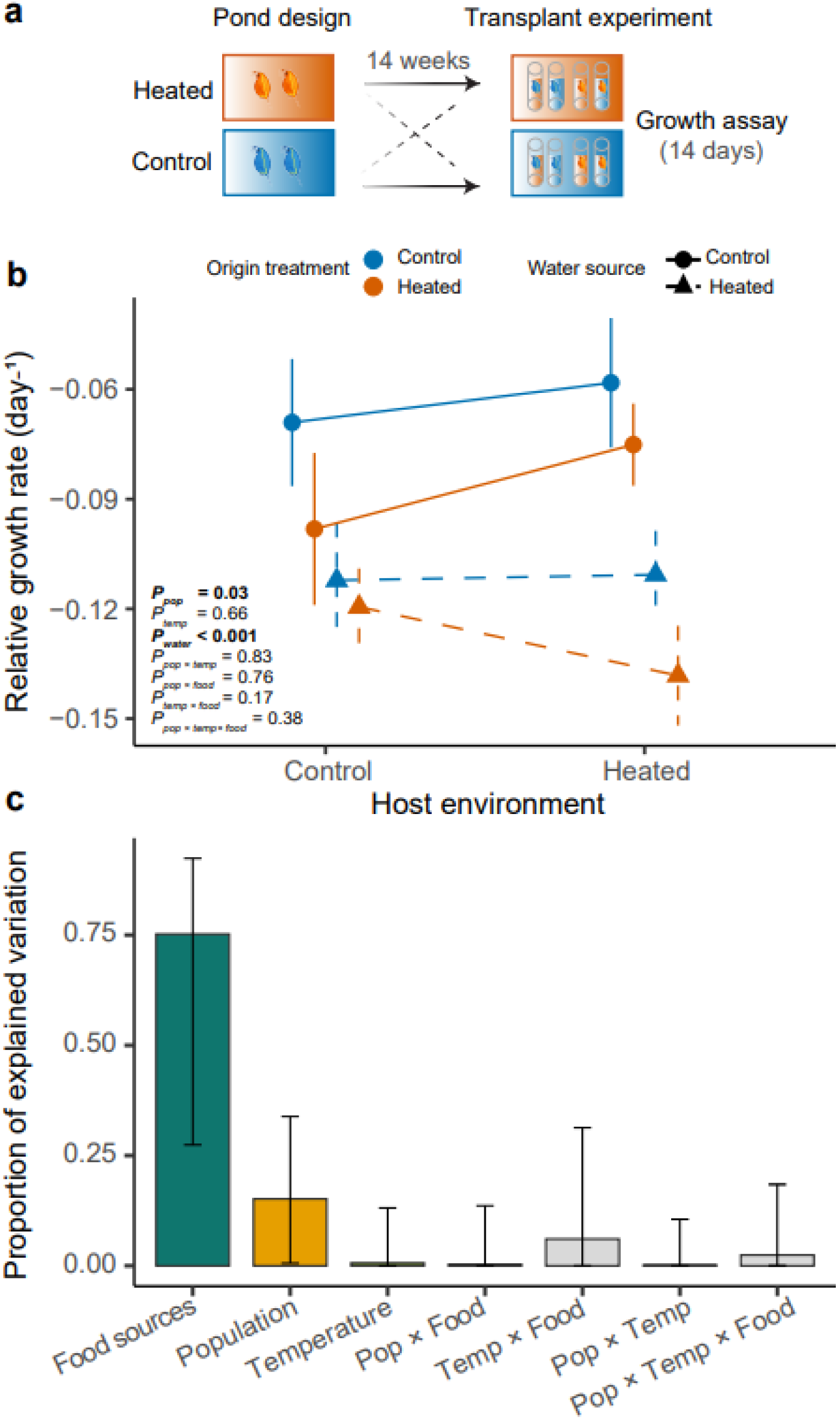
Water-mediated environmental conditions (food sources) dominate variation in *D. magna* population growth. **a,** Schematic of the reciprocal transplant experiment. Populations originating from control and heated mesocosms were reciprocally transplanted after 14 weeks of experimental warming into either control or heated host temperature treatments. Transplants were performed using either water from the recipient pond or water from the alternative treatment to separate direct temperature effects from water-mediated environmental influences. Population growth was quantified over a 14-day assay. **b,** Relative growth rate of *D. magna* across population origin (control, blue; heated, orange), host temperature (control vs. heated), and water source (solid lines indicate control water; dashed lines indicate heated water). Points represent means and bars indicate ± SE. P-values are derived from a linear mixed-effects model including population origin, host temperature, and water source as fixed effects and pond and block as random effects. **c,** Proportional contribution of each fixed effect and interaction to variation in RGR of *D. magna*, based on Type III sums of squares from the fitted linear mixed-effects model. Bars represent the proportion of explained variance attributed to each term; error bars indicate 95% confidence intervals.

*Daphnia* growth rate was affected primarily by water source rather than on host temperature. *Daphnia* exposed to water from warmed ponds grew less than those maintained in control water, regardless of their population origin or recipient thermal environment (*P* < 0.001; Fig. 5b). Population origin had a smaller but significant effect on growth (*P* = 0.03), whereas host temperature had no detectable effect (*P* = 0.66). Consistent with this result, water source accounted for 75.21% of the explained variation in relative growth rate, compared with 15.18% for population origin and 0.65% for host temperature (Fig. 5c).

These results indicate that reduced *Daphnia* performance under warming was associated mainly with changes in pond conditions rather than with direct temperature increase during the transplant assay.

## Discussion

By combining replicated outdoor warming with a naturally occurring heatwave, we show that near-future warming can become ecologically consequential, reorganizing a multitrophic freshwater community across trophic levels. Warmed ponds showed a shift towards duckweed dominance, reduced snail abundance, increased aphid populations, lower phytoplankton biomass, and reduced *Daphnia* abundance. This is especially the case when an extreme event pushes temperatures across a consumer-relevant thermal threshold. The thermal vulnerability of *L. stagnalis* during the heatwave provides a likely entry point for this cascade: during the July 2025 heatwave, temperatures even in the middle of the water column in warmed ponds exceeded the reported sublethal thermal stress threshold of *L. stagnalis* (Hoefnagel & Verberk 2017), whereas control ponds remained below this threshold. This pattern indicates a marked loss of cool deep-water refuge for a benthic grazer during a short extreme event in our mesocosms. Although the strong warming at depth might have been accentuated by the bottom-heating design of our warming system, the underlying mechanism is consistent with patterns observed in nature: subsurface and bottom heatwaves occur in lakes, are increasing in frequency, and can reduce the availability of thermal refuge (Woolway *et al*. 2025). Shallow freshwater systems may be particularly vulnerable because they often provide only weak vertical thermal buffering.

The thermal vulnerability of *L. stagnalis* during the heatwave, a keystone organism in the community, provides a likely mechanism linking warming to the observed community changes. Shortly after the heatwave, snail abundance was very low in warmed ponds, and the climate-chamber experiment showed that herbivory peaked at intermediate temperatures but dropped sharply at 32 °C, where mortality also increased. This reduction in grazing pressure was associated with increased duckweed dominance under warming, consistent with the negative relationship between snail abundance and duckweed cover across ponds. Although warming itself could also enhance duckweed growth (Fig. 4), this effect was less pronounced in populations that had experienced selection within the community. Increased duckweed dominance was in turn associated with broader changes in community structure, likely through reduced light availability (Fig. S2) and altered pond conditions, contributing to lower algae abundance and reduced *Daphnia* performance.

Reduced Daphnia performance was best explained by water source rather than host temperature, suggesting that warming altered pond conditions indirectly. This likely reflected reduced algal food availability in warmed ponds, consistent with lower chlorophyll-a concentrations and with previous mesocosm work showing that duckweed dominance can suppress algal biomass and *Daphnia* abundance (Schäfer *et al*. 2025). Interestingly, the drop in *Daphnia* abundance preceded the major increase in duckweed coverage in the heated ponds. This is likely due to higher abundance of algae (reflected in higher chlorophyll-a concentrations) in heated ponds, giving *Daphnia* an advantage. Then *Daphnia* grew faster and overgrazed the algae, which led to their population crash by week 3 in the heated ponds. *Daphnia* abundance followed with a 2-week lag, declining by week 5 as food availability became limiting in the heated ponds. The heatwave occurred as this decline was already underway. Because the recorded water temperatures are within the thermal ranges of *Daphnia* survival and reproduction (28–35 °C and 26–32 °C respectively), the heatwave thus mostly likely contributed to the observed decline in *Daphnia* populations through indirect food-web effects (Seefeldt & Ebert 2019). The contrasting responses of different trophic groups highlight the importance of incorporating species interactions and community structure at a high temporal resolution when predicting ecological responses to future climate warming.

Field-realistic warming of aquatic communities has historically required electrically powered heating elements and automated control systems, which have enabled foundational insights and are more precise but can limit scalability (Yvon-Durocher *et al*. 2010; Stewart *et al*. 2013). Alternative approaches include open-top chambers, which passively trap heat without electrical infrastructure, but have typically been applied to smaller water bodies with temperature measurements restricted to shallow surface layers (Netten *et al*. 2008). Transplantation of aquatic communities across altitudinal gradients has also been used to exploit natural temperature differences (Johnsen *et al*. 2020), though this approach is constrained by limited site replication and logistical feasibility.

Our solar-powered approach reduces operational energy requirements while preserving ambient temporal variability, an important advantage because many warming experiments still rely on static mean offsets that inadequately capture ecologically consequential variability and extreme events (Thompson *et al*. 2013; Vasseur *et al*. 2014). Because organisms’ responses depend on non-linear thermal performance curves, preserving such variability allows us to realistically evaluate whether warming increases or decreases performance, particularly during naturally occurring extreme conditions. Because the warming system is inexpensive to install, adopting this system at different regions can provide new mechanistic insights into communities’ responses to warming at a global scale.

## Materials and Methods

### Experimental setup

The outdoor mesocosm experiment was conducted at the Botanical Garden in Mainz, Germany, from May to October 2025 and consisted of 32 experimental ponds. Approximately six weeks before the start of the experiment, all ponds (2.4 m diameter, 1.2 m depth; pond volume ∼5.4 m^3^) were drained completely and cleaned with a high-pressure cleaner. Each pond contained two perforated Euroboxes (60 cm × 40 cm × 32 cm; volume 64 L) filled with mixed leaf litter collected from the botanical garden to provide nutrients and habitat for aquatic organisms.

We used a blocked design in which ponds were grouped into 16 blocks of two ponds each. Within each block, one pond was assigned to the heated treatment and the other to the control treatment, resulting in 16 ponds per treatment. The orientation of the treatments was alternated between each row to ensure an equal distribution of treatments across the area and to reduce positional effects. Both heated and control ponds were equipped with the same solar-powered infrastructure (Tables S1 and S2), and ponds within the same block shared the same heating unit. Water flow from the solar panels was regulated by a switching valve that continuously compared pond temperature with the temperature of water exiting the panels. When panel water was cooler (e.g. during night, early morning, or cold periods), the valve interrupted circulation to prevent cooling of the heated ponds. Under these conditions, residual flow could slightly cool the control ponds, thereby maintaining a temperature difference even when solar heating was inactive.

To assemble the experimental plant population, we used 96 *Spirodela polyrhiza* genotypes originating from four geographic groups: America, Europe, India, and Southeast Asia. Four weeks before the start of the experiment, genotypes were surface-sterilized with sodium hypochlorite and cefotaxime, and then cultivated under controlled conditions (26 °C, 16 h light : 8 h dark, 135 µmol m^-2^ s^-1^) in N-medium (Appenroth *et al*. 1996).

Ponds were filled with tap water one week before the start of the experiment. One day before experimental initiation, water was mixed among all 32 ponds via pumps to reduce initial environmental differences. The experiment began on 21 May 2025 (week 0). Each pond received 20 fronds of each *S. polyrhiza* genotype (1,920 fronds in total), which were introduced directly into the pond and constituted a free-floating mixed population. In addition, 75 waterlily aphids (*Rhopalosiphum nymphaeae*; 15 genotypes, five individuals per genotype), approximately 1,200 *Daphnia magna* individuals from a mixture of 122 genotypes were introduced into each pond. The *Daphnia* were the same mix of genotypes used in the earlier experiment (Schäfer *et al*. 2025). Furthermore, eight *Lymnaea stagnalis* individuals from an in-house culture at Eawag (Dübendorf, Switzerland) were added to each pond. To promote zooplankton establishment, ponds also received 450 mL of a mixed algal culture (*Tetradesmus* spp. and *Nannochloropsis* spp.) previously used to feed *D. magna* in the laboratory. KH_2_PO_4_ (12.3 g) was added to each pond to enrich phosphorus availability. Ponds were covered with a fine-mesh net (120 µm mesh opening) that allowed light penetration and duckweed growth while reducing aphid dispersal and limiting access by larger animals.

### Floating grid system

To maintain individual duckweed genotypes under field conditions, we additionally established a floating grid system made of Styrofoam (60 cm × 60 cm × 2 cm) positioned in two corners of each pond. The system consisted of four grids, each accommodating 25 genotypes, allowing all 96 *S. polyrhiza* genotypes to be maintained individually. Each grid contained cups that were open at the bottom, allowing exchange with the surrounding pond water while keeping genotypes physically separated. This setup allowed us to expose genotypes to pond-specific abiotic and biotic conditions while preserving their genetic identity for later reconstruction experiments.

### Population dynamics in the experimental ponds

Community dynamics were monitored every two weeks throughout the 2025 growing season (n = 16 ponds per treatment).

To assess the overall effects of the heated treatment on the *S. polyrhiza* populations, we estimated duckweed surface coverage (mixed population) in each pond through visual inspections conducted throughout the experiment. These estimates were cross-validated using photographs taken at each timepoint with a GoPro Hero 9 camera, similar to our previous work (Schäfer *et al*. 2025). To assess phenotypic differences between populations at week 9 of the mesocosm experiment, approximately 500 fronds were collected from each mixed population to quantify dry biomass and surface area per frond.

Aphid density was estimated from close-up photographs taken with a mobile phone camera from six different positions per pond. In each image, fronds and associated aphids were manually counted, and aphid density was expressed as the number of aphids per frond. To estimate total aphid abundance at the community level, mean aphid density per pond was multiplied by duckweed coverage for each pond and timepoint.

We quantified *D. magna* abundance in the experimental ponds by sampling vertical water-column profiles from four locations per pond using a Leibold sampler (a 140-cm plastic tube, 6 cm in diameter, closable at the bottom with a wire). A total of 9 L of water was collected in a bucket per pond. From each sample, two subsamples were taken: 50 mL for nutrient analyses and 1 L stored in dark bottles for chlorophyll-*a* analysis. The remaining water was filtered through a 120-µm mesh, and retained individuals were preserved in ≥98 % ethanol and counted manually in the laboratory.

Snail abundance was quantified through visual inspections of the pond walls, and individuals were counted manually. Initially (week 0), 8 individuals were introduced into each pond. From week 7 onwards, snail density was assessed by counting individuals within a standardised 1/12 section of the pond side wall, spanning from the surface to the bottom of the pond.

Chlorophyll-*a* concentrations, used as a proxy for phytoplankton abundance, were assessed from pigment extracts using HPLC–PDA. From each 1-L subsample, water was filtered sequentially onto 25 mm GF/F glass fibre filters (Fisherbrand) using a water pump. The filters were removed from the holder, gently dried on tissue paper, placed in cryovials, and stored at –20 °C until further processing. Prior to chromatographic analysis, filters were lyophilized under dark conditions and pigments were extracted in 1.3 mL of solvent (81.1 % methanol, 10.8 % ethyl acetate, 8.1 % water and 180 mmol/L ammonium acetate). Extraction was performed by bead beating (BeadBugTM 3 microtube Homogenizer, Benchmark Scientific, USA; 3 × 30 s at 400 rpm) with intermittent cooling on ice, using a glass bead mixture (0.25–0.50 mm beads filling 500 µL of the tube, with four additional 4.0 ± 0.3 mm beads added). The supernatant was transferred to a new 1.5 mL tube. Extracts were centrifuged at 31,550 × g and 4 °C for 1 min. Subsequently, 300 µL of the supernatant was mixed with 30 µL of water, and 100 µL was injected into the HPLC system for analysis as previously described (Schäfer *et al*. 2025).

### Nutrients and abiotic environment in the experimental ponds

To test for differences in water chemistry between heated and control ponds, nutrient concentrations were analysed by ion chromatography using a Metrohm Eco IC system equipped with an 863 Compact IC autosampler (Metrohm, Switzerland).

At each sampling time point (every two weeks), water was collected from four different locations of the pond covering the full water-column profile (as described above). A total of 10 mL per pond was collected for each anion analysis. Anions were separated using a Metrosep A supp 19 Guard/4.0 pre-column, eluent 8.0 mmol L^-1^ Na_2_CO_3_, 0.25 mmol L^-1^ NaHCO_3_, and 5 % acetonitrile, flow rate 0.7 mL min^-1^, and an injection volume of 10 µL. Anion quantification was performed using external calibration with five-point standard curves (Cl⁻ and SO ^2^⁻: 200, 100, 50, 25, and 12.5 mg L^-1^; NO ⁻ : 1, 0.5, 0.25, 0.125, and 0.063 mg L^-1^; PO ^3^⁻: 20, 10, 5, 2.5, and 1.25 mg L^-1^).

In parallel, we also measured pH, temperature, dissolved oxygen, and conductivity using dipping probes (Multi 3630 IDS with an FDO 925 oxygen sensor, a SenTix 940 pH sensor, and a TetraCon 925 conductivity sensor) submerged at a depth of 65 cm below the water level.

### Thermal monitoring and heatwave

Water temperature (°C) and light intensity (lux) were recorded in each pond using temperature loggers (HOBO MX2202) installed at two depths: the water surface (0 cm) and 65 cm below the surface. Loggers recorded temperature and light at 10-min intervals during the season. Raw measurements were aggregated to daily summaries for each pond and depth, from which daily mean and maximum temperatures and light were calculated. To confirm that mesocosm conditions tracked natural environmental variation, daily mean pond temperatures were compared with ambient air temperatures recorded by external loggers placed outside the experimental ponds.

A documented regional heatwave occurred between 28 June–02 July 2025 in Mainz, Western Germany (https://opendata.dwd.de/climate_environment/CDC/), and this real-world climatic event was used to define heatwave conditions in the experiment.

To quantify treatment effect size, daily temperature differences (ΔT) were calculated as ΔT = T_heated_ – T_control_. Median ΔT values were calculated separately for each depth using both daily mean and maximum temperatures, providing a robust estimate of treatment-induced warming independent of short-term variability.

### Reciprocal transplant experiments

To test whether reduced *D. magna* performance under warming was driven by direct temperature effects or by warming-induced environmental change, we conducted a reciprocal transplant assay during week 14 of the mesocosm experiment (01 September 2025). Individuals were collected from six vertical positions within each pond using a 140-cm PVC tube spanning the water column. From these collections, 200 similarly sized adult females without eggs were selected and evenly assigned to four transplant combinations (n = 50 per combination). Individuals were transplanted either into another pond of the same treatment or into a pond of the alternative treatment. To separate direct thermal effects from water-mediated environmental effects, each transplant was performed using either water from the recipient pond or water from the pond of the alternative treatment. The water was filtered through a 120 µm mesh beforehand to exclude other zooplankton. Assays were conducted in vertical PVC tubes (115 cm long, 6 cm diameter), closed at the bottom and open at the top, for 14 days. At the end of the assay, individuals were collected and counted immediately. Relative growth rate (RGR) was calculated as (ln(N_end_+1) - ln(N_start_+1))/ΔTime, where N_end_ and N_start_ represent the number of individuals at the end and beginning of the assay, respectively, and ΔTime represents the duration of the assay (days). A constant of 1 was added prior to log transformation to avoid undefined values for zero counts.

For duckweed reciprocal transplant assays, mixed populations originating from heated and control ponds were transplanted either into another pond of the same treatment or into a pond of the alternative treatment. Fronds were collected by sampling 80 spots from distinct areas of each pond, pooled, and mixed, and 130 fronds were subsequently used to initiate each assay. Growth assays were performed in floating boxes (17 cm × 12 cm × 11.5 cm) fitted with a 0.6-mm mesh on the bottom. After two weeks, fronds were collected and counted to calculate RGR.

To further distinguish evolutionary responses from phenotypic plasticity, we reconstructed the initial genotype composition using individual genotypes growing in the floating grid. The initial (reconstructed) populations comprised 65 of the original 96 genotypes, as the remaining genotypes were absent or present in insufficient numbers to establish the assay at standardised starting frequencies (two fronds per genotype). Populations were reassembled from warming and control treatment origins and transplanted following the same reciprocal design described above, with populations placed either into the same treatment or into the alternative treatment. The assay lasted for 14 days, after which fronds were collected and counted to calculate RGR.

### Climate-chamber assay of temperature effects on plant growth and herbivory

To isolate direct temperature effects from other environmental factors, we performed a climate-chamber microcosm experiment across three independent rounds. In each round, mixed duckweed populations were established with two fronds per genotype, resulting in 190 fronds per microcosm (95 genotypes; one genotype was not viable in the laboratory collection).

Experimental units were maintained in eight replicate boxes (16 cm × 13 cm × 6 cm) per temperature treatment, each containing 500 mL of homogenised pond water. Water was collected from at least 20 ponds. Within each round, collected water was filtered through a 120 µm mesh, pooled and thoroughly homogenised prior to allocation among experimental boxes.

Temperature regimes were derived from field measurements recorded during the peak growing season. Four experimental temperature treatments (20 °C, 24 °C, 28 °C, and 32 °C) were selected to represent the observed thermal range across treatments and depths, ensuring ecologically realistic yet controlled conditions.

Within each round, temperature treatments were applied sequentially across four consecutive days. For each temperature, eight replicate microcosms were established, five assigned to the herbivory treatment and three serving as growth controls without snails to account for intrinsic plant growth during the 22-hour feeding period. In herbivory treatments, three *L. stagnalis* individuals (size 22–24 mm) were introduced per replicate.

Prior to the herbivory assays, snails were starved and acclimated to their respective experimental temperature for 24 hours. Both growth and susceptibility assays lasted 22 hours. Fronds were photographed and counted to quantify plant growth and the number of consumed fronds. Snail mortality was also recorded, and temperature-specific mortality rates were calculated.

### Statistical analyses

The impact of the heated treatment on population dynamics (duckweed surface coverage, snail abundance, aphid density, *Daphnia* abundance, and chlorophyll-*a* concentrations; n = 16 per treatment) was analysed using linear mixed-effects models. Treatment (heated or control) and sampling timepoint were included as fixed effects, and pond and block were included as random effects. Block was included as a random effect because pond pairs shared the same solar-powered heating unit. Models were specified as response ∼ treatment × timepoint + (1|pond) + (1|block). Significance of fixed effects was assessed using F-tests with denominator degrees of freedom estimated using the Kenward-Roger approximation.

The effect of warming on duckweed morphology (biomass and area per frond) was tested using linear mixed-effects models. Treatment (heated or control) was included as a fixed effect, and pond and block were included as random effects.

To disentangle the effects of population origin, host temperature, and water-mediated environmental conditions on *D. magna* population growth, we conducted a reciprocal transplant experiment and calculated the relative growth rate (RGR) over the 14-day assay period. We fitted a fully factorial linear mixed-effects model including population origin, host temperature, and water source as fixed effects, with pond and block included as random effects. Fixed effects were evaluated using Type III sums of squares, with each term expressed as a proportion of total explained variation. Uncertainty in proportional contributions was estimated using a pond-level cluster bootstrap procedure, in which recipient ponds were resampled with replacement and the model refitted for each bootstrap replicate. 95% confidence intervals were derived from the empirical bootstrap distribution.

To assess evolutionary and environmentally induced plastic responses in duckweed, we conducted a transplant assay using evolved and initial populations. RGR was calculated over the 14-day assay. We tested the effects of elevated temperature on growth in mixed populations (evolved) by fitting a linear mixed-effects model specifying population and host temperature (heated vs. control) as fixed effects, and pond and block as random effects. The effect of plasticity on growth was assessed in the initial populations using the same specifications in the model.

To separate the direct effects of temperature from other biotic and environmental factors, we conducted an additional indoor climate-chamber experiment under controlled conditions. Temperature effects on plant growth and herbivore consumption were analysed using linear mixed-effects models. For each response variable (growth rate and control-corrected consumption rate per snail), temperature (20, 24, 28, and 32 °C) was included as a fixed effect, and experimental round was included as a random effect. RGR was calculated as the change in frond number over the 22-hour assay. Herbivore consumption was calculated as the difference between mean frond change in control microcosms and frond change in herbivory treatments, divided by the average number of snails present during the assay. Snail mortality was calculated for each temperature within each experimental round as the proportion of individuals that died during the 22-hour feeding period (number of deaths divided by the initial number of snails per temperature), and mean mortality rates were calculated across rounds.

## Acknowledgments

We thank the Botanical Garden of the University of Mainz for their support in establishing and maintaining the pond facility and for assistance with water collection. We also thank Pascal Blaschke, Julia Doskoc, Annira Klink, Anna Sofia Kolehmainen, Sian Stockton, and Thoomke Roth for assistance with the pond experiments, frond counting, and duckweed cultivation and Christoph Vorburger from Swiss Federal Institute of Aquatic Science and Technology for providing the snails. We thank the workshop at the University of Mainz for constructing equipment such as the floating grids, the samplers, mesh cover supports, and PVC-tubes for the *Daphnia* transplant assay. This work is supported by European Research Council under the European Union‘s Horizon 2020 research and innovation program (Grant agreement No. 101125029 to S.X.).

