## Supplementary Information for "Near-future warming amplifies natural heatwave impacts and reorganizes freshwater communities"

**Figures**

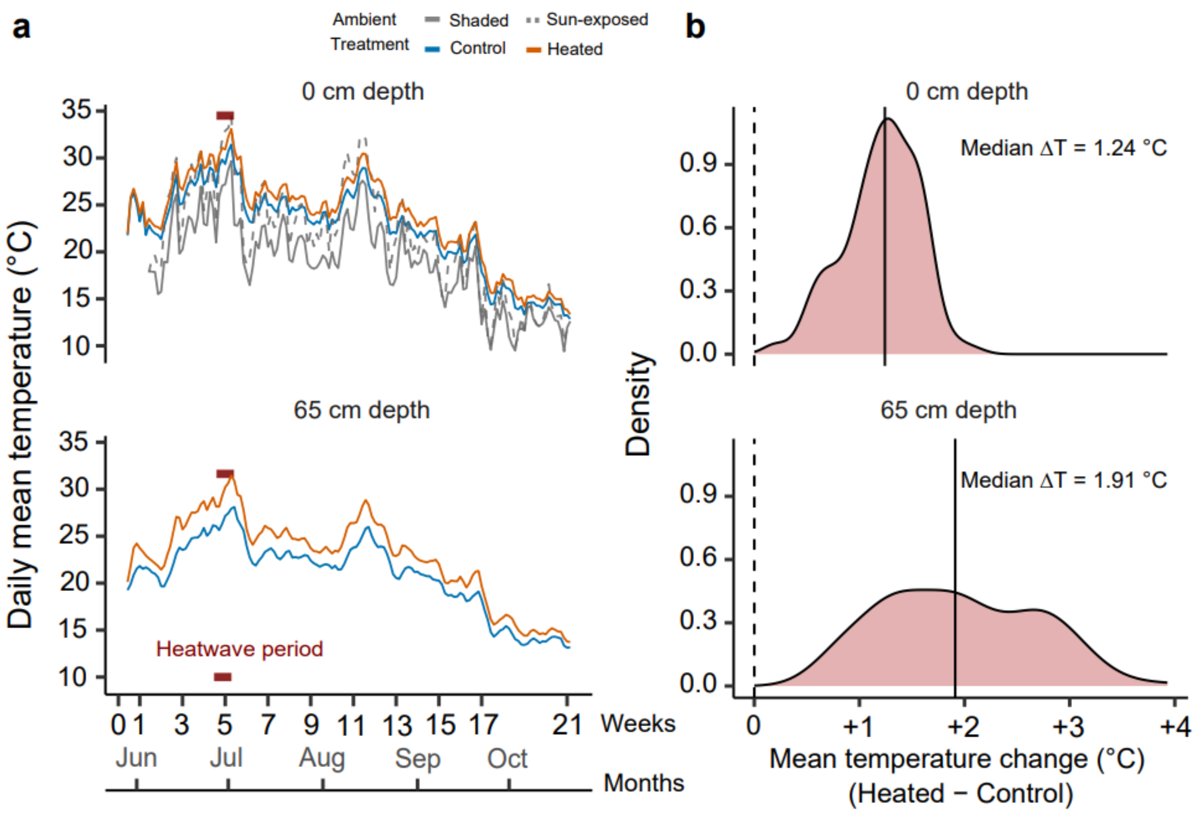


**Fig. S1. Warming altered thermal regimes across depths and amplified temperature differences during the growing season. a,** Daily mean temperature at 0 cm and 65 cm depth in heated (orange) and control (blue) ponds, shown alongside ambient temperatures from shaded (solid grey) and sun-exposed (dashed grey) external loggers. The heatwave period is indicated in red colour. **b,** Distribution of daily temperature differences (heated – control), with vertical lines indicating median $\Delta T$.

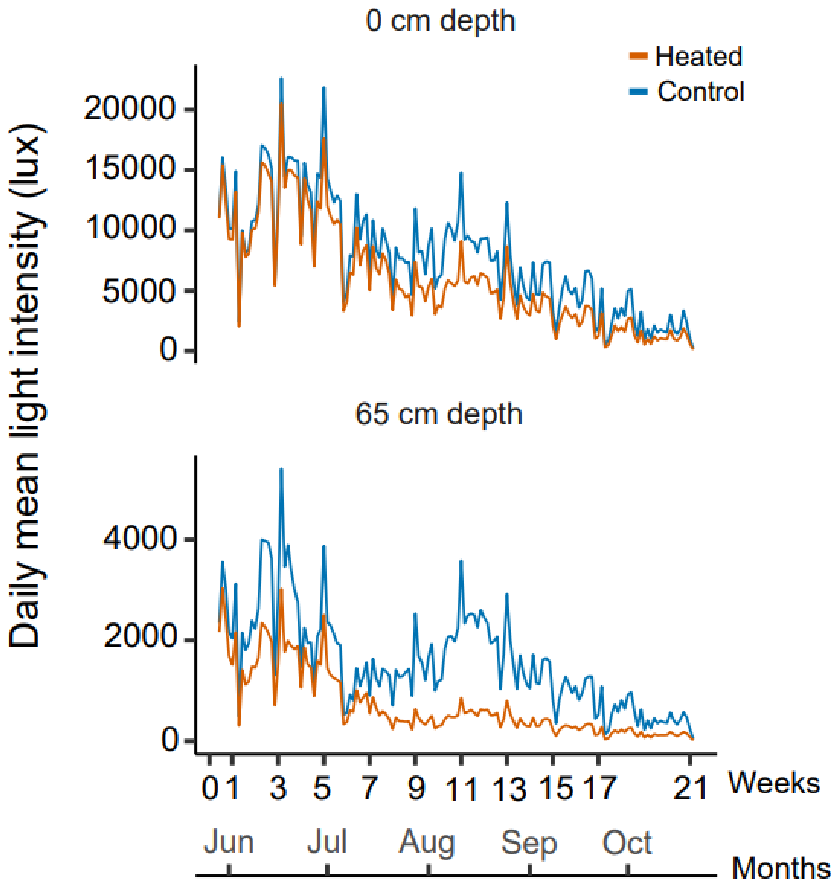


**Fig. S2. Reduced light availability at depth in heated ponds**. Daily mean light intensity (lux) at 0 cm and 65 cm depth in heated (orange) and control (blue) ponds across the 2025 growing season.

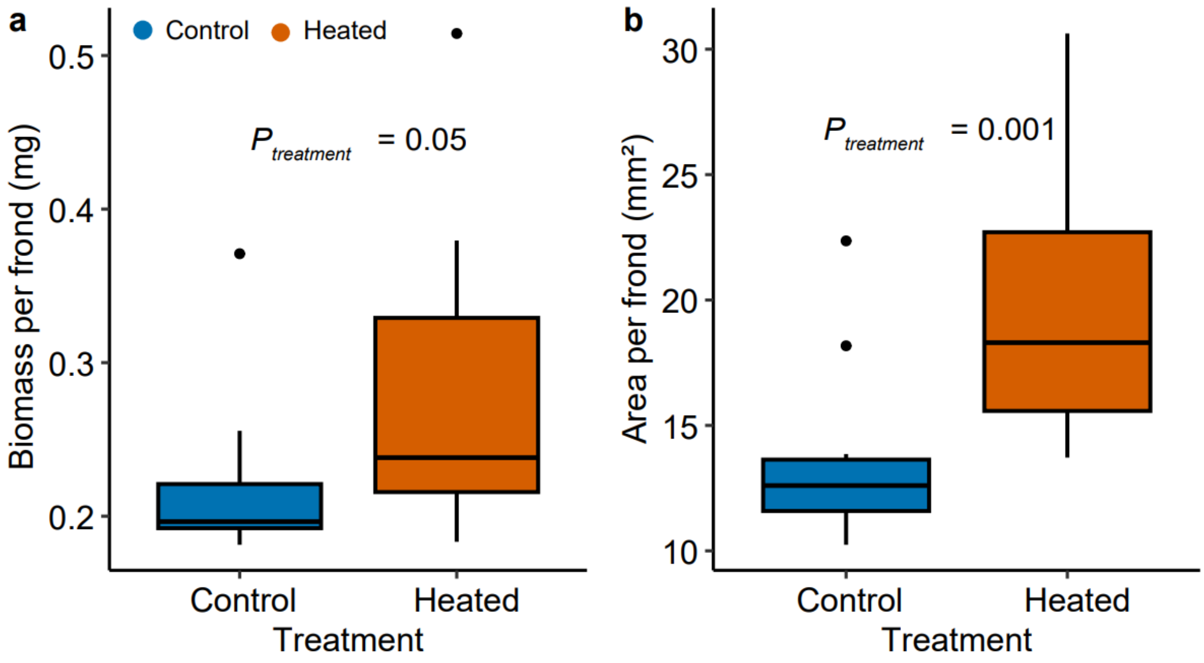


**Fig. S3. Duckweed fronds were larger and heavier under warming. a,** Dry biomass per frond (mg) and **b,** frond area ($mm^2$) measured from 500 undamaged fronds from *S. polyrhiza* populations exposed to heated (orange) and control (blue). P-values indicate treatment effects from linear mixed-effects models (n = 16 per treatment).

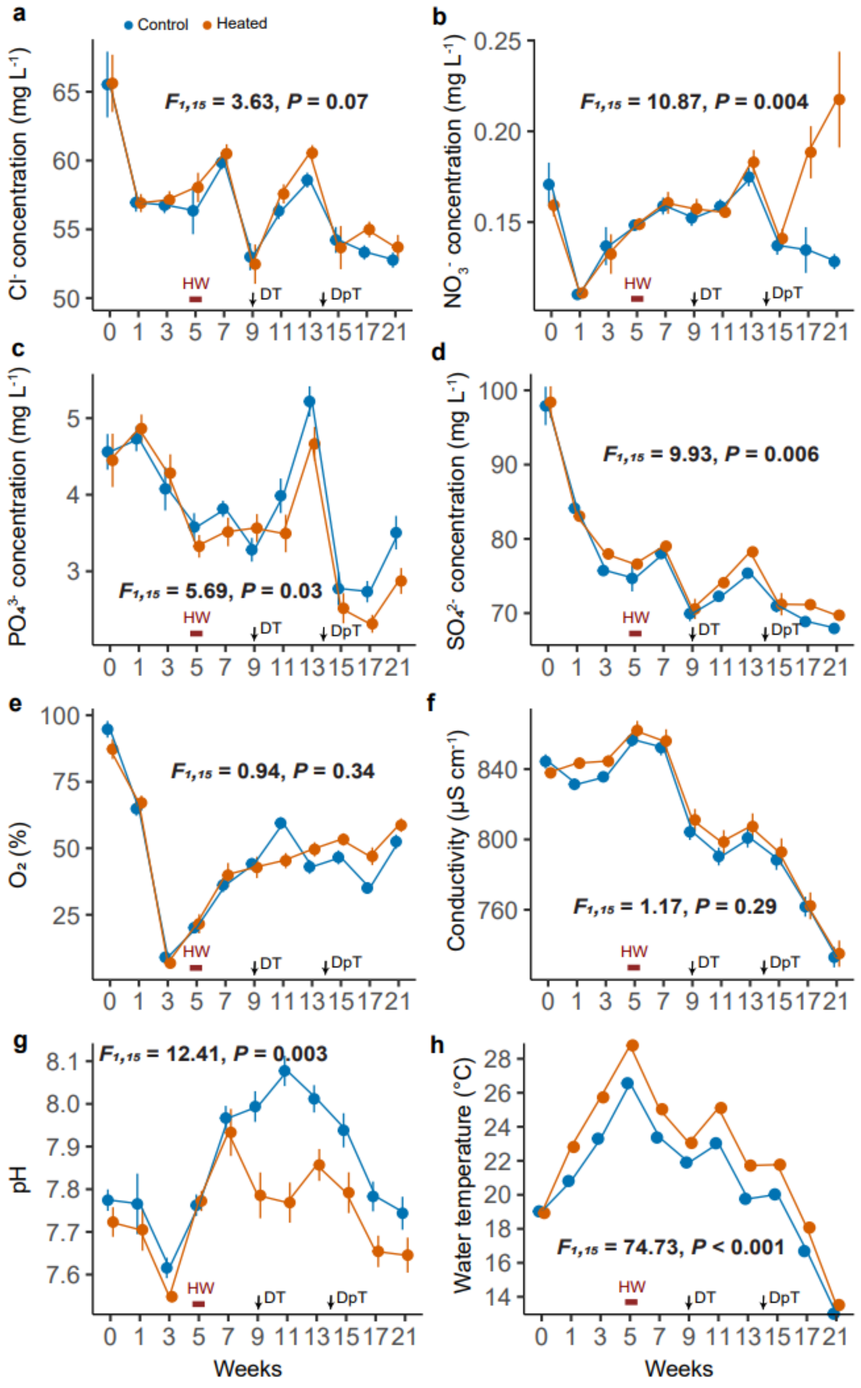


**Fig. S4. Temporal dynamics of nutrient concentrations and physicochemical variables in control and heated ponds during the 2025 growing season. a,** Chloride. **b,** Nitrate. **c,** Phosphate. **d,** Sulfate. **e,** Oxygen. **f,** Conductivity. **g,** pH. **h,** Water temperature at 65 cm depth; from the dipping sonde. Orange and blue lines indicate heated and control treatments, respectively. Red horizontal bar indicates the heatwave period (28 June–02 July 2025). DT, duckweed transplant timepoint; DpT, *Daphnia* transplant timepoint. The x-axis shows experimental duration in weeks. Values represent means ± SE. Statistics are from linear mixed-effects models with treatment and timepoint as fixed effects and pond and block as random effects.

**Tables**

**Table S1 Core infrastructure costs of the outdoor mesocosm experiment.** Costs include materials required to construct a 32-pond outdoor mesocosm system. Most components represent one-time investments and are reusable across experimental runs.

| Component | Details | Cost (€) | Reusable |
|---|---|---|---|
| Ponds | 32 ponds | 43,495 | Yes |
| Insulation materials | Bubble wraps (Regular and aluminium), plastic sheets, aluminium protection band | 1,303.61 | Yes |
| Mesh | Pond mesh covers | 3,200 | Yes |
| Mesh fixation materials | Metal clamps, rubber strings | 940.50 | Yes |
| Stones | ~1200 kg | 435 | Yes |
| Solar panels | 32 panels | 2,546.62 | Yes |
| Pumps | 16 units (40 W each) | 1,552.80 | Yes |
| Pump buckets | | 100.48 | Yes |
| Temperature control valves | 16 (+3 replacement units) | 3,781 | Yes |
| Tubing and connectors | | 1,088.34 | Yes |
| Electrical components | Sockets | 224 | Yes |
| Miscellaneous materials | String, duct tape, cable ties, Teflon tape | 542.44 | Mixed |
| **Total** | | **59,209.79** | |
| **Total per pond** | | **1,850.32** | |

**Table S2. Annual operational energy costs of the mesocosm system.** Energy consumption is generated by pumps (640 W total for 32 ponds), resulting in very low annual energy demand.

| Component | Details | Cost (€. $yr^{-1}$) |
|---|---|---|
| Energy consumption | 16 pumps (640 W total): ~ 5606.4 kWh. $yr^{-1}$ | 1,400-1,700 |
| Temperature control system | Energy for valves and sensors | Negligible |
| Battery HOBO loggers | Battery replacement (CR2032; 1-2 per logger per year) | ~ 25-50 |
| **Total** | | **1,425-1,750** |
| **Total per pond** | | **45-55** |

**Table S3. Effect of warming and sampling timepoint on community dynamics during the 2025 growing season.** Results are from linear mixed-effects models with treatment and timepoint as fixed effects, and pond and block as a random effect. Values shown are fixed-effect estimates ± SE.

| | **Factor** | **Estimate ± SE** | $df_{res}$ | $F$ | $p$ |
|---|---|---|---|---|---|
| Duckweed coverage | Treatment | 1.07 ± 9.20 | 15 | 14.38 | **0.001** |
| | Timepoint | 1.29 ± 0.32 | 254 | 123.09 | **< 0.001** |
| | Treatment × Timepoint | 2.48 ± 0.46 | 254 | 29.58 | **< 0.001** |
| Snail abundance | Treatment | -36.20 ± 14.78 | 15 | 32.47 | **< 0.001** |
| | Timepoint | 2.70 ± 0.60 | 190 | 22.38 | **< 0.001** |
| | Treatment × Timepoint | -1.40 ± 0.85 | 190 | 2.74 | 0.09 |
| Aphid per frond | Treatment | 0.11 ± 0.14 | 15 | 4.54 | **0.05** |
| | Timepoint | -0.065 ± 0.008 | 237 | 129.63 | **< 0.001** |
| | Treatment × Timepoint | 0.002 ± 0.011 | 237 | 0.03 | 0.87 |
| *Daphnia* abundance | Treatment | -194.33 ± 70.70 | 15 | 21.76 | **< 0.001** |
| | Timepoint | -20.76 ± 3.68 | 254 | 61.39 | **< 0.001** |
| | Treatment × Timepoint | 0.75 ± 5.20 | 254 | 0.02 | 0.88 |
| Chlorophyll-*a* concentration | Treatment | -1.03 ± 0.36 | 15 | 33.33 | **< 0.001** |
| | Timepoint | -0.081 ± 0.02 | 252 | 33.81 | **< 0.001** |
| | Treatment × Timepoint | -0.0007 ± 0.02 | 252 | 0.001 | 0.98 |

**Table S4. Effects of experimental warming and sampling timepoint on water nutrient concentrations and physicochemical parameters during the 2025 growing season.** Linear mixed-effects models were used to test the effects of warming, sampling timepoint, and their interaction on water nutrient concentrations and physicochemical parameters in experimental ponds (n = 16 ponds per treatment). Treatment (heated vs. control) and timepoint were included as fixed effects, and pond and block as random effects.

| **Response** | **Factor** | **Estimate ± SE** | $df_{res}$ | $F$ | $p$ |
|---|---|---|---|---|---|
| Chloride ($Cl^-$) | Treatment | 0.63 ± 0.98 | 15 | 3.63 | 0.07 |
| | Timepoint | -0.24 ± 0.06 | 254 | 36.09 | **< 0.001** |
| | Treatment × Timepoint | 0.01 ± 0.07 | 254 | 0.05 | 0.82 |
| Nitrate ($NO_3^-$) | Treatment | -0.03 ± 0.01 | 15 | 10.87 | **0.004** |
| | Timepoint | -0.001 ± 0.0006 | 254 | 7.86 | **0.005** |
| | Treatment × Timepoint | 0.004 ± 0.0008 | 254 | 30.46 | **< 0.001** |
| Phosphate ($PO_4^{3-}$) | Treatment | 0.13 ± 0.25 | 15 | 5.69 | **0.03** |
| | Timepoint | -0.03 ± 0.01 | 254 | 29.45 | **< 0.001** |
| | Treatment × Timepoint | -0.03 ± 0.02 | 254 | 3.12 | 0.07 |
| Sulfate ($SO_4^{2-}$) | Treatment | 1.63 ± 1.06 | 15 | 9.93 | **0.006** |
| | Timepoint | -0.45 ± 0.05 | 254 | 120.47 | **< 0.001** |
| | Treatment × Timepoint | 0.001 ± 0.08 | 254 | 0.0002 | 0.99 |
| Dissolved oxygen (%) | Treatment | -4.65 ± 3.95 | 15 | 0.94 | 0.34 |
| | Timepoint | 1.83 ± 0.21 | 254 | 211.01 | **< 0.001** |
| | Treatment × Timepoint | 0.60 ± 0.29 | 254 | 4.20 | **0.04** |
| Conductivity (µS/cm) | Treatment | 8.78 ± 6.27 | 15 | 1.17 | 0.29 |
| | Timepoint | -6.50 ± 0.26 | 254 | 1364.60 | **< 0.001** |
| | Treatment × Timepoint | -0.32 ± 0.36 | 254 | 0.79 | 0.37 |
| pH | Treatment | -0.07± 0.05 | 15 | 12.41 | **0.003** |
| | Timepoint | 0.002 ± 0.002 | 254 | 0.06 | 0.81 |
| | Treatment × Timepoint | -0.005 ± 0.003 | 254 | 1.77 | 0.18 |
| Temperature (°C) | Treatment | 2.53 ± 0.44 | 15 | 74.73 | **< 0.001** |
| | Timepoint | -0.63 ± 0.02 | 254 | 1463.65 | **< 0.001** |
| | Treatment × Timepoint | -0.07 ± 0.03 | 254 | 4.61 | **0.03** |
